# Dissecting the genetic basis of DUS characters in Pakistani wheat varieties

**DOI:** 10.64898/2026.11.01.763533

**Authors:** Ihtisham Ul Haq, Ijaz Ahmed Qureshi, Muhammad Ans Hussain, Rana Arsalan Javaid, Fawad Sheikh, Zeeshan Abbas, Muhammad Azam

## Abstract

Distinctness, Uniformity and Stability (DUS) testing is fundamental to varietal characterization and the protection of Plant Breeders’ Rights (PBRs), but conventional DUS assessment relies predominantly on morphological characteristics whose expression may be influenced by environmental conditions, developmental stage and observer-related variation. Integrating character-specific molecular markers with conventional DUS assessment may therefore improve the reproducibility and genetic resolution of varietal identification. This study investigated the genetic architecture of DUS characteristics in a panel of 102 historical and modern Pakistani wheat varieties using genome-wide association studies (GWAS). Phenotypic variation was evaluated using distribution, correlation and hierarchical clustering analyses, while the germplasm was genotyped using a wheat genotyping-by-targeted sequencing platform. After quality filtering, 21,670 high-quality single-nucleotide polymorphisms (SNPs) were retained for analyses of genetic diversity, population structure, linkage disequilibrium (LD) and marker-trait associations. Population structure analysis indicated extensive admixture, with cross-entropy supporting K = 9 ancestral components, whereas hierarchical genetic clustering separated the germplasm into five broad groups. The first three principal components explained 8.09%, 7.20% and 5.39% of the total genetic variation, respectively. GWAS using BLINK and FarmCPU identified significant associations for several diagnostically important DUS characteristics. Growth habit was strongly associated with SNP 1B_129176704, with an additional signal on chromosome 5B. Shattering showed its strongest association at 1D_445256324, accompanied by several closely linked SNPs in the same region and a secondary significant locus at 7B_703502819. Flag leaf length was associated with a cluster of SNPs on chromosome 6A, while tiller density showed a major association with 5D_205488820. These findings demonstrate substantial phenotypic and genetic diversity within Pakistani wheat germplasm and identify genomic regions that merit validation as character-specific molecular tools to complement conventional DUS testing and PBR implementation.

## INTRODUCTION

Food insecurity remains a major global challenge and is increasingly intensified by climate-related stresses. United Nations Sustainable Development Goal 2 aims to end hunger, achieve food security, improve nutrition and promote sustainable agriculture by 2030 (UN, 2015). The development and deployment of climate-resilient crop varieties can contribute to this objective; however, plant breeding requires substantial long-term investment. Effective systems for protecting new plant varieties are therefore important for encouraging continued innovation and supporting returns on breeding investment. The International Union for the Protection of New Varieties of Plants (UPOV) Convention provides an international framework for the protection of new plant varieties (UPOV, 1991). In Pakistan, the Plant Breeders’ Rights Act, 2016 (GoP, 2016) and the Plant Breeders’ Rights Rules, 2018 (GoP, 2018) provide the national legal framework for Plant Breeders’ Rights (PBRs), broadly following the principles of the UPOV Convention 1991.

Distinctness, Uniformity and Stability (DUS) testing is central to the grant of PBRs and is conventionally based on the expression of qualitative, pseudo-qualitative and quantitative morphological characteristics across prescribed testing conditions and seasons. Because phenotypic expression may be influenced by environmental conditions, developmental stage and scoring subjectivity, character-specific molecular information could provide an additional layer of evidence for varietal characterization. The UPOV Technical Working Group on Biochemical and Molecular Techniques and DNA Profiling has considered approaches in which molecular markers linked to specific characteristics can support DUS testing (UPOV, 2020). More recently, genome-wide association analysis has been used to identify loci controlling DUS characteristics in wheat (Zanella *et al*., 2026), and genomic prediction has been proposed as a complementary strategy for managing DUS reference collections and supporting trait-specific distinctness decisions (Roberts *et al*., 2026).

Wheat is one of the world’s most important food crops and contributes substantially to global calorie and protein intake (Erenstein *et al*., 2022). Its broad adaptation, storability and nutritional importance make it central to global food security (Iqbal *et al*., 2022). Wheat is also of particular importance to Pakistan, where it occupies a major proportion of the cultivated area and contributes substantially to national food supply (GoP, 2025). Genome-wide association studies conducted in Pakistani and related wheat germplasm have identified significant loci for several agronomic and adaptive traits. Under heat stress, significant marker-trait associations have been reported for kernel-weight reduction (Hussain *et al*., 2026). Allelic and haplotypic variation at *TaHST1* has been associated with heat tolerance (Khan *et al*., 2022), while loci and haplotypes related to reduced plant height have been investigated in relation to *Rht* genes (Suleman *et al*., 2025). Additional studies have reported candidate genes related to glucose metabolism, ion transport, hormone responses and starch synthesis (Zhou and Irshad, 2026), as well as quantitative trait nucleotides associated with canopy architecture, flag leaf area and flag leaf width (Farhan *et al*., 2026).

The present study evaluated DUS characteristics in 102 Pakistani wheat varieties representing historical and modern germplasm. The primary objective was to identify SNP associations for DUS characteristics using high-throughput GWAS models and to assess whether robust marker-trait associations could provide a basis for future character-specific molecular tools. Such tools are intended to complement, rather than replace, conventional DUS phenotyping and may ultimately strengthen varietal identification and PBR enforcement in Pakistan.

## MATERIALS AND METHODS

### Plant material and DUS phenotyping

Identity records and DUS phenotypic data for 102 Pakistani wheat varieties, including historical lines and modern cultivars, were obtained from the DUS Office, Federal Seed Certification and Registration Department (FSC&RD), Ministry of National Food Security and Research, Islamabad, Pakistan. Qualitative and pseudo-qualitative characteristics were converted to numerical DUS notes for analysis, as described in Supplementary Table S1.

Phenotypic data were analysed in R using the packages dplyr, tidyr, ggplot2, dendextend and corrplot. Trait distributions were visualized using violin plots, while hierarchical clustering was used to evaluate phenotypic similarity among varieties. Correlation analysis was performed to examine the direction and magnitude of relationships among characteristics. Where traits were measured on different scales, standardized *z*-scores were used to facilitate direct comparison.

### Genotyping and genetic analyses

The wheat panel was genotyped using a 16K wheat genotyping-by-targeted sequencing (GBTS) platform (Rasheed and Fayyaz, 2023). The initial dataset contained approximately 37,000 SNPs. Quality filtering was performed using BCFtools, and markers with a minor allele frequency (MAF) < 0.05 were removed. After filtering, 21,670 high-quality SNPs were retained for downstream analyses.

Linkage disequilibrium (LD) was evaluated as pairwise marker correlation (*r*²) using PLINK 2.0. LD decay was assessed by plotting *r*² against physical distance in the Chinese Spring wheat reference genome (Consortium *et al*., 2018) for the A, B and D subgenomes. An *r*² threshold of 0.20 was used as a reference for LD decay. Principal component analysis (PCA) was performed on the filtered SNP dataset, and the first three principal components were visualized in three-dimensional space.

Population structure was evaluated using a model-based ancestry framework implemented in the LEA package in R across *K* = 2-10 ancestral components. Model fit was compared using cross-entropy with the optimal value of *K* selected from the minimum cross-entropy. A hierarchical dendrogram based on genome-wide marker distances was also generated. Together, PCA, ancestry analysis and hierarchical clustering were used to assess relatedness, genetic differentiation and the potential influence of population structure on association analysis.

### Genome-wide association studies and post-GWAS analysis

Genome-wide association analyses were conducted using Bayesian-information and Linkage-disequilibrium Iteratively Nested Keyway (BLINK) and Fixed and random model Circulating Probability Unification (FarmCPU) implemented in GAPIT in R. For each characteristic, raw *P* values were adjusted using the Benjamini-Hochberg false discovery rate (BH-FDR) at α = 0.05 and the Bonferroni family-wise error threshold at α = 0.05. With 21,670 markers, the nominal Bonferroni threshold was approximately *P* = 2.31 × 10⁻⁶. Manhattan and quantile-quantile (QQ) plots were examined to evaluate the distribution of association signals and model calibration. Genomic inflation factors were evaluated from the QQ-plot outputs.

Significant SNPs identified by GWAS were positioned relative to the IWGSC Chinese Spring reference genome (Consortium *et al*., 2018). Candidate genes were considered within LD-defined intervals surrounding significant SNPs. Candidate regions were interpreted with reference to gene position and published functional information. To evaluate the phenotypic effects of significant marker-trait associations, post-GWAS genotype-class comparisons were performed using appropriate parametric or non-parametric statistical procedures following the general approach described by (Nawade *et al*., 2025). Kruskal-Wallis tests were used for the genotype-class comparisons reported for the principal loci.

## RESULTS

### Phenotypic variation and trait relationships

The evaluated wheat varieties showed substantial phenotypic variation across the DUS characteristics. Standardization to *z*-scores enabled direct comparison among traits measured on different scales. The violin plots revealed marked differences among characteristics in distribution width, central tendency and skewness, indicating substantial trait-specific variation within the germplasm. Qualitative and pseudo-qualitative characteristics generally showed more discrete or concentrated distributions, as expected from their note-based scoring system.

Hierarchical clustering further demonstrated phenotypic differentiation among the varieties. Based on their combined standardized trait profiles, the genotypes were separated into five major phenotypic clusters (Figure 1). Two clusters contained a relatively large proportion of the germplasm, whereas the remaining groups contained smaller and more distinct sets of varieties. The separation of these groups at comparatively large linkage distances indicates meaningful differences in overall phenotypic profiles.

**Figure 1.**
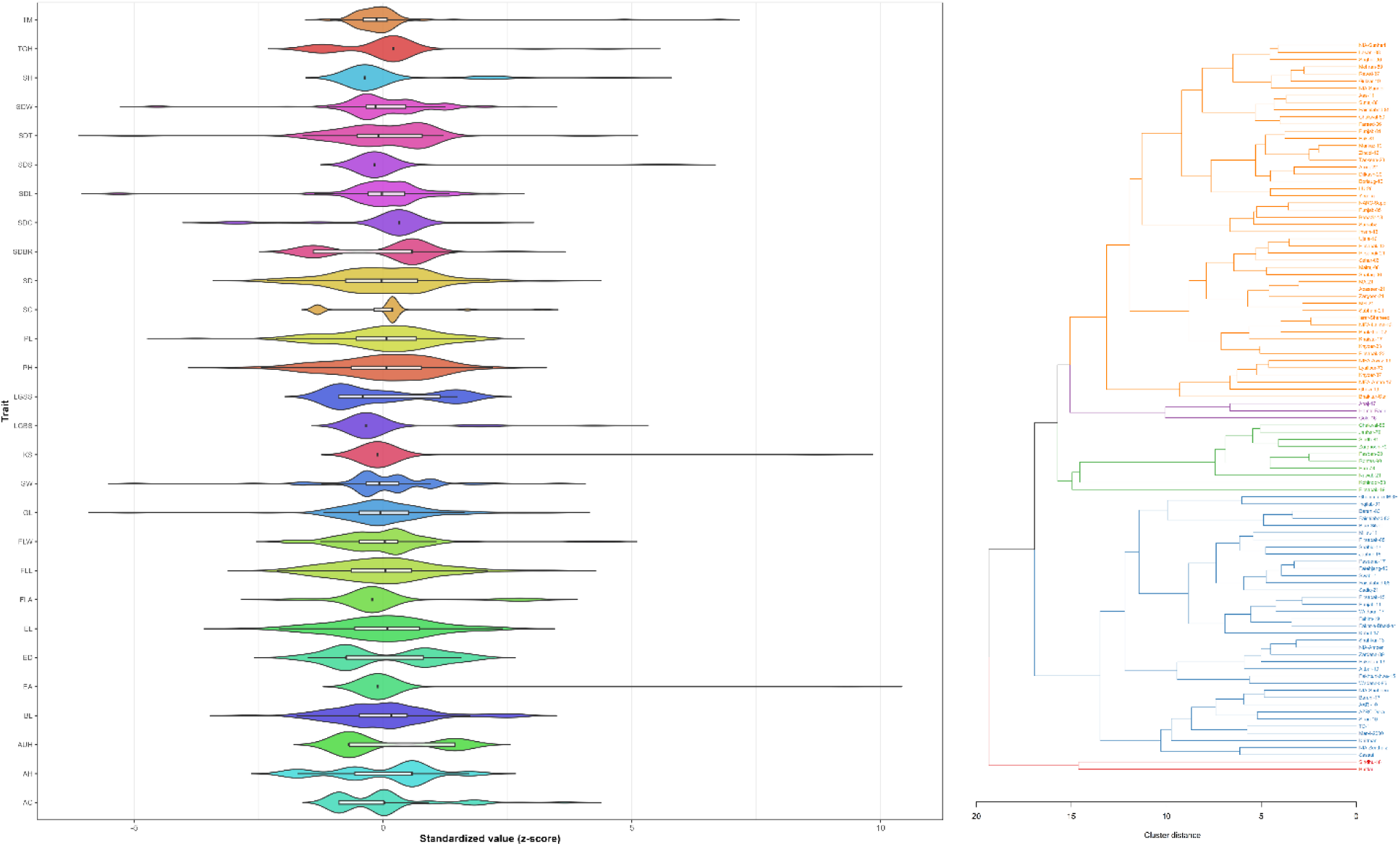
Phenotypic variation and hierarchical clustering of DUS characteristics in Pakistani wheat varieties. Distribution of standardized phenotypic values and hierarchical relationships among the 102 wheat varieties evaluated for DUS characteristics. The violin plots (left) show the distribution, density and central tendency of individual characteristics after standardization to *z*-scores. The hierarchical dendrogram (right) was generated from the combined phenotypic profiles and separates the evaluated varieties into five major phenotypic clusters, illustrating the extent of morphological diversity within the germplasm panel.

The correlation matrix showed both positive and negative relationships among traits, although most pairwise correlations were relatively weak (Figure 2). This pattern indicates that many DUS characteristics capture largely independent dimensions of varietal morphology. Collectively, the distribution, clustering and correlation analyses demonstrate broad phenotypic diversity within the panel and provide an appropriate phenotypic basis for subsequent association analysis.

**Figure 2.**
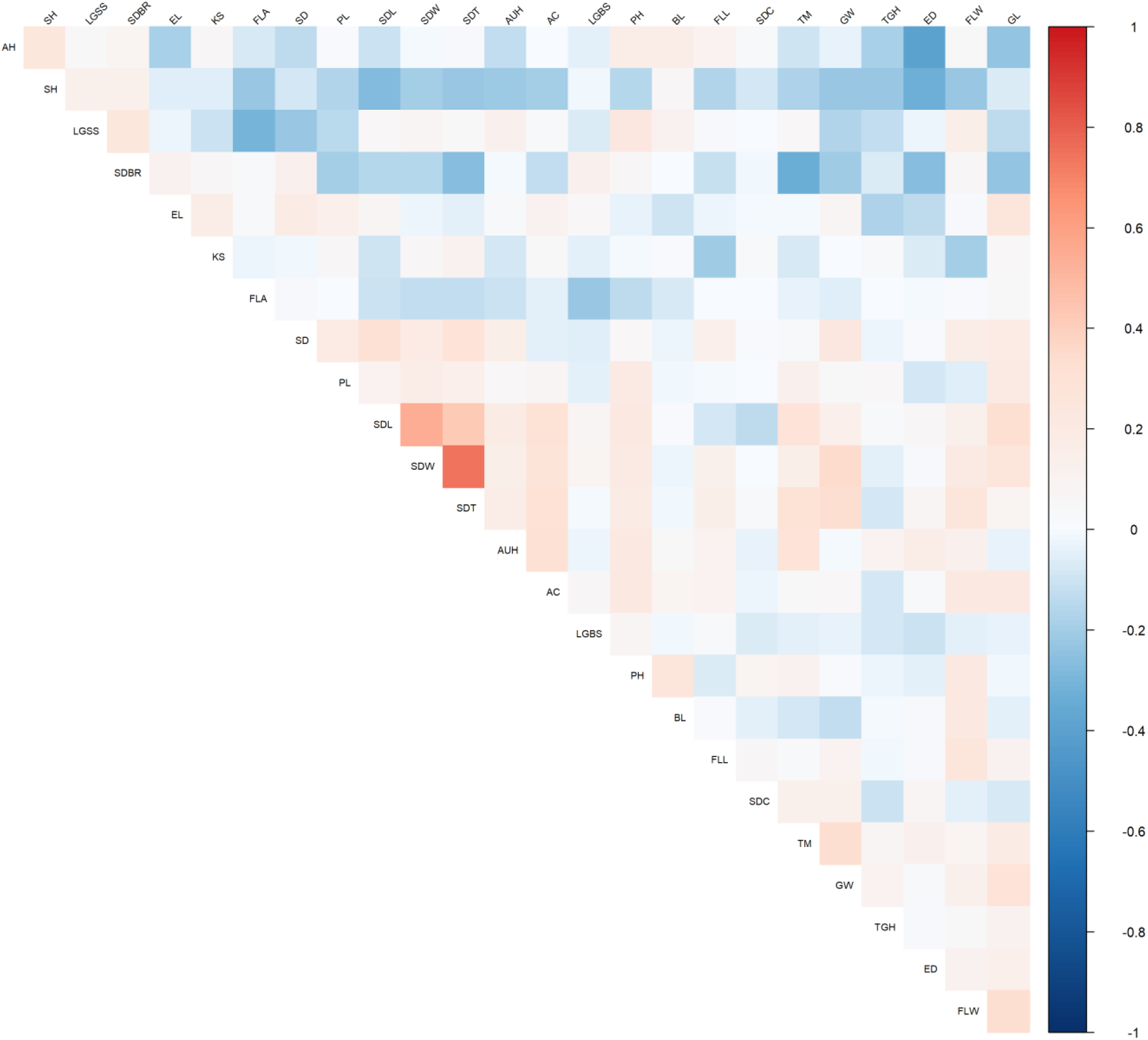
Correlation matrix of DUS characteristics evaluated in Pakistani wheat varieties. Pairwise correlation coefficients among the evaluated DUS characteristics are presented as a lower-triangular heatmap. The colour scale ranges from −1 to +1, with red indicating positive correlations, blue indicating negative correlations and values approaching white indicating weak or negligible relationships.

### Genotyping and genetic analyses

Model-based population structure was evaluated across K = 2-10 ancestral components. At all values of K, most accessions contained contributions from more than one ancestry component, indicating extensive admixture among the evaluated wheat varieties. Cross-entropy decreased from approximately 0.59 at K = 2 to a minimum of approximately 0.518 at K = 9, followed by a slight increase to approximately 0.524 at K = 10 (Figure 3). Accordingly, K = 9 was selected as the best-supported model for population structure. The progressive reduction in cross-entropy up to K = 9 indicates that the panel contains substantial substructure that is not adequately represented by only a few ancestral groups. At K = 9, several sets of accessions showed relatively high contributions from specific ancestry components, while extensive admixture remained evident, consistent with historical gene flow and shared breeding backgrounds.

**Figure 3.**
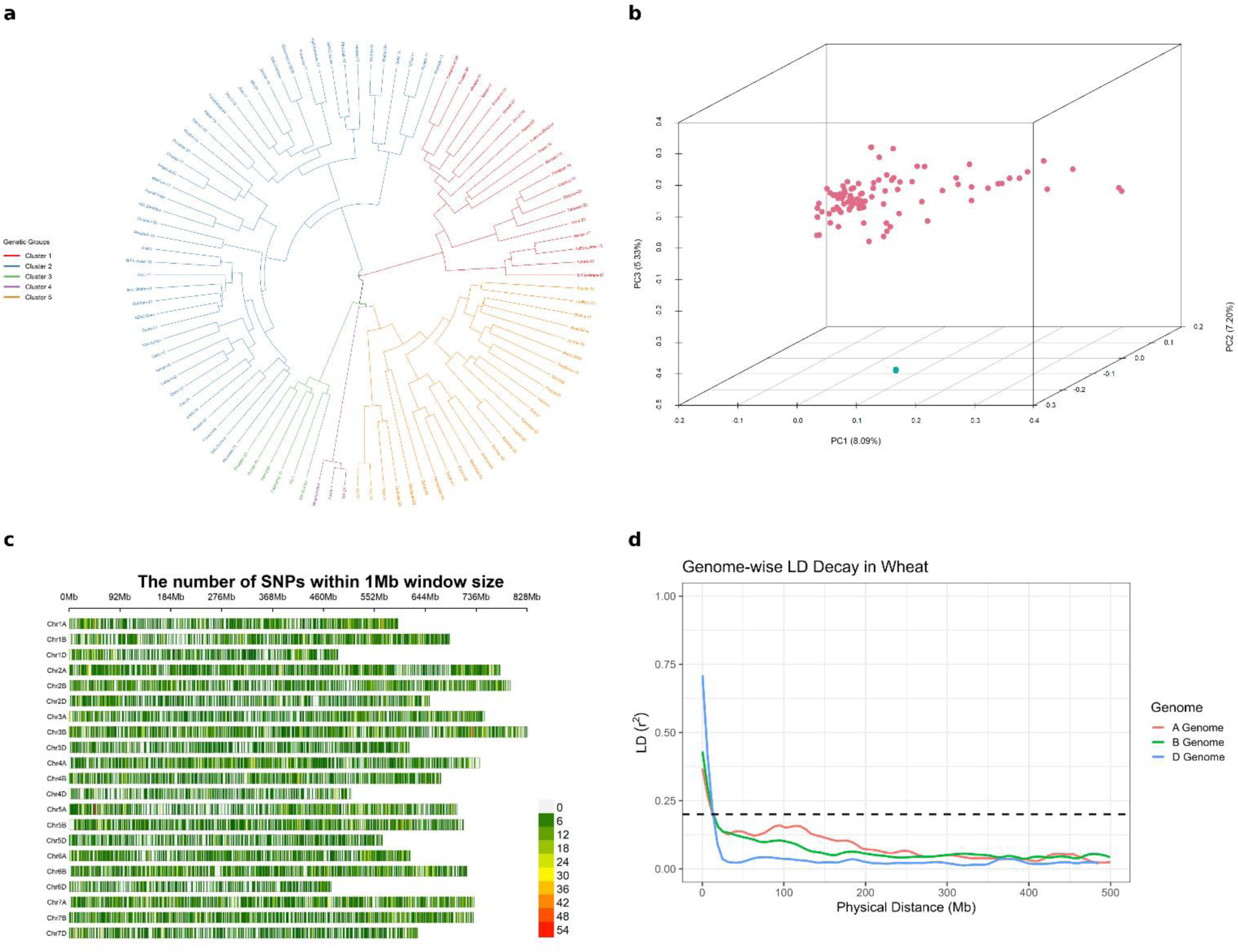
Genome-wide genetic diversity, principal component analysis, SNP distribution and linkage disequilibrium decay in Pakistani wheat varieties. **(a)** Circular hierarchical dendrogram based on genome-wide SNP data, showing the genetic relationships among the 102 wheat varieties and their separation into five broad genetic clusters. **(b)** Three-dimensional principal component analysis (PCA) based on the filtered SNP dataset; PC1, PC2 and PC3 explain 8.09%, 7.20% and 5.39% of the total genetic variation, respectively. **(c)** Genome-wide distribution of SNPs across the 21 wheat chromosomes, expressed as the number of markers within 1-Mb windows. **(d)** Linkage disequilibrium (LD) decay for the A, B and D subgenomes, expressed as pairwise *r*² against physical distance (Mb). The horizontal dashed line represents the reference LD threshold of *r*² = 0.20.

The circular hierarchical dendrogram provided complementary evidence of genetic differentiation. Based on genome-wide marker distances, the accessions were broadly separated into five major genetic clusters. One cluster contained a substantial proportion of the accessions and showed considerable internal branching, whereas the smaller clusters comprised more closely related sets of genotypes (Figure 4). The five broad dendrogram groups and nine ancestry components represent different levels of population structure rather than conflicting results. Hierarchical clustering summarizes broad genome-wide relatedness, whereas the model-based ancestry analysis resolves finer substructure and admixture within the major groups. Together, these analyses indicate a hierarchical genetic organization within the Pakistani wheat panel.

**Figure 4.**
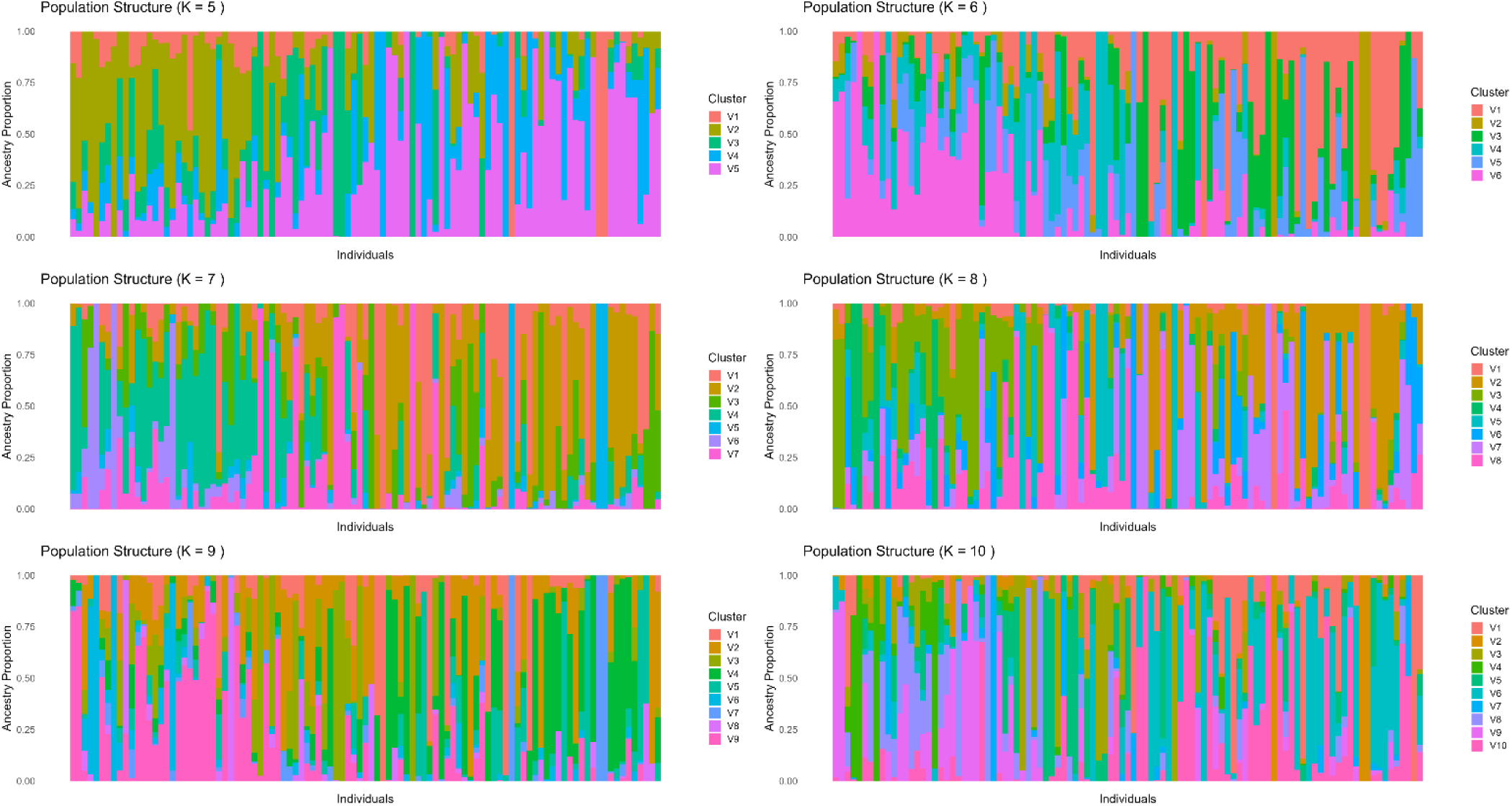
Model-based population structure of Pakistani wheat varieties across different numbers of ancestral populations. Ancestry proportions of the 102 wheat varieties estimated for K = 5–10 ancestral components. Each vertical bar represents an individual variety and different coloured segments indicate the estimated proportional contribution of each ancestral component. Cross-entropy analysis identified K = 9 as the best-supported population-structure model.

The first three principal components explained 8.09%, 7.20% and 5.39% of the total genetic variation for PC1, PC2 and PC3, respectively. Most accessions occupied a relatively compact region of the three-dimensional PCA space, consistent with shared ancestry and genetic exchange among breeding materials. Nevertheless, differentiation was evident along all three axes, and at least one accession was clearly separated from the main group. The moderate proportion of variance explained by each component suggests that population differentiation is distributed across several dimensions rather than being dominated by a single major axis.

Genome-wide SNP-density analysis showed marker coverage across all 21 wheat chromosomes of the A, B and D subgenomes. SNP abundance, assessed in 1-Mb windows, varied among chromosomes and genomic regions. Some intervals contained relatively few SNPs, whereas others contained more than 50 markers per 1-Mb window. Despite this heterogeneity, the filtered marker set provided broad genome-wide coverage for population and association analyses. LD, expressed as pairwise *r*², declined with increasing physical distance in all three subgenomes, although the decay trajectories differed. The D genome showed comparatively high LD at very short distances followed by a relatively rapid decline, whereas the A and B genomes showed more gradual decay and retained higher LD over longer distances. These subgenome-specific patterns indicate differences in the extent of allelic association across the wheat genome.

Overall, the genetic analyses demonstrated substantial diversity, pronounced admixture and hierarchical population structure in the study panel. The combined ancestry, clustering and PCA results support the use of models that account for population structure when interpreting marker-trait associations.

### Genome-wide association studies

GWAS for growth habit (TGH) identified two principal loci, with the strongest and most consistent signal on chromosome 1B and a second signal on chromosome 5B. In both BLINK and FarmCPU, the associated regions showed clear departures from the genomic background. The QQ plots showed that deviation from the expected null distribution was concentrated primarily in the upper tail, consistent with a limited number of true association signals rather than widespread inflation. The lead SNP on chromosome 1B, 1B_129176704, was highly significant (P = 3.756 × 10⁻⁹; BH-FDR = 8.140 × 10⁻⁵; Bonferroni-adjusted P = 8.140 × 10⁻⁵). The locus had a positive allelic effect of 1.610 and a low MAF of 0.053, indicating that the effect allele was relatively uncommon in the study panel (Figure 5). A second marker, 5B_586820792, was also associated with growth habit (P = 2.333 × 10⁻⁶; BH-FDR = 2.528 × 10⁻²). Its Bonferroni-adjusted P value of 0.05056 placed the association at the boundary of the strict family-wise significance criterion. Genotype-class comparisons supported phenotypic differentiation at both loci. For 1B_129176704, the Kruskal-Wallis test was significant (P = 0.00878), and the A/A class showed the highest growth-habit scores, the G/G class showed intermediate values and the G/A class showed the lowest values. For 5B_586820792, genotype classes also differed significantly (Kruskal-Wallis P = 0.00376), with the C/C genotype showing higher scores than the T/T and T/C classes (Figure 6). These results identify chromosome 1B as the major growth-habit region in this panel, with additional evidence for a secondary locus on chromosome 5B.

**Figure 5.**
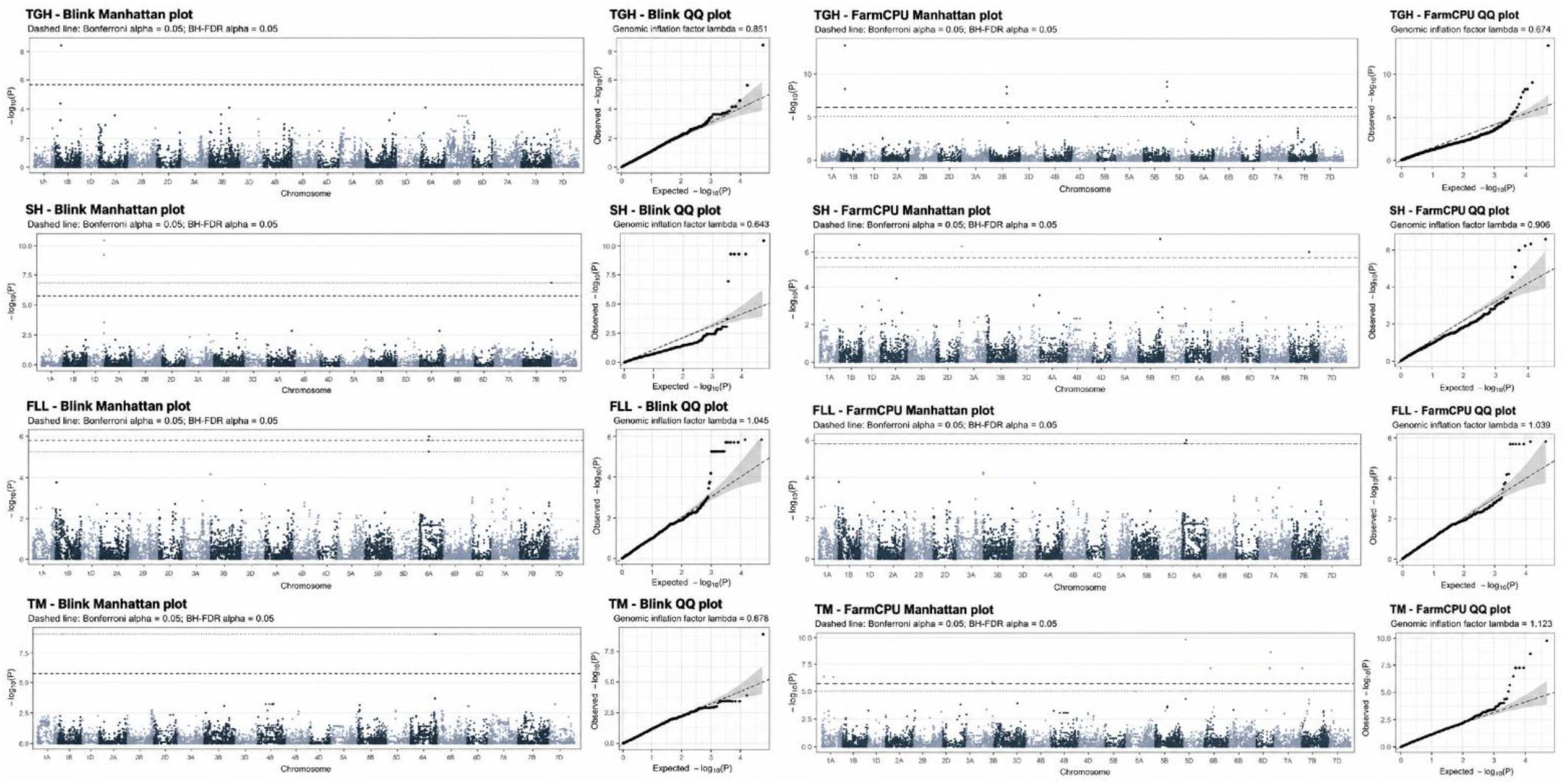
Genome-wide association analysis of growth habit, shattering, flag leaf length and tiller density using BLINK and FarmCPU models. Manhattan and quantile–quantile (QQ) plots generated for growth habit (TGH), shattering (SH), flag leaf length (FLL) and tiller density (TM) using the BLINK and FarmCPU GWAS models. For each trait, the BLINK Manhattan and QQ plots are shown alongside the corresponding FarmCPU Manhattan and QQ plots. Manhattan plots display marker significance as −log₁₀(*P*) across the 21 wheat chromosomes. Horizontal dashed lines indicate the Bonferroni and Benjamini–Hochberg false discovery rate (BH-FDR) significance thresholds at α = 0.05. QQ plots compare observed and expected −log₁₀(*P*) values, with genomic inflation factors (λ) shown for assessment of model calibration and residual population-structure effects.

**Figure 6.**
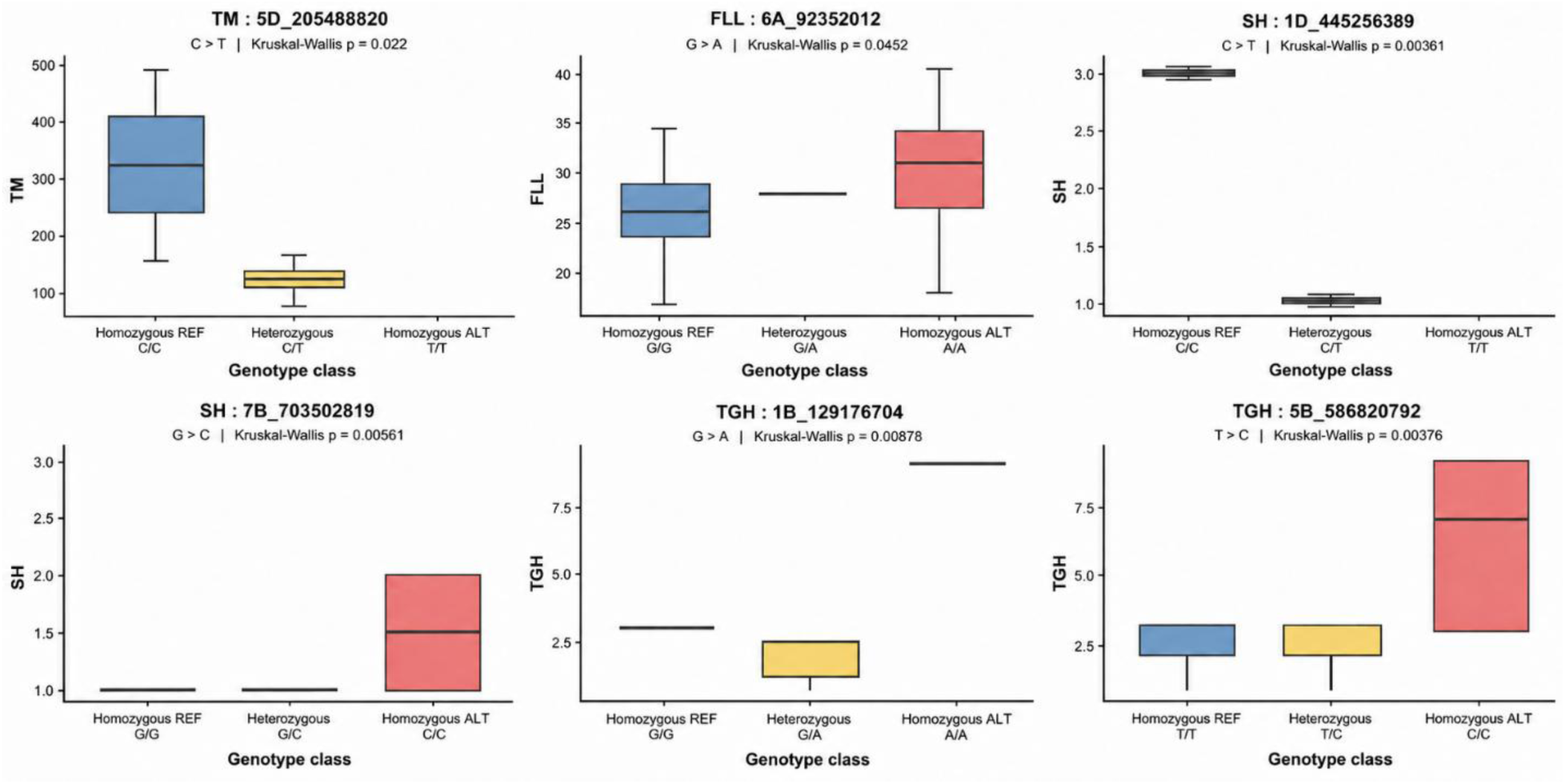
Genotype-specific phenotypic effects of representative GWAS-associated loci for tiller density, flag leaf length, shattering and growth habit in Pakistani wheat varieties. Boxplots show the phenotypic distributions of genotype classes for six significant or representative marker–trait associations: **(a)** tiller density (TM) at 5D_205488820, **(b)** flag leaf length (FLL) at 6A_92352012, **(c)** shattering (SH) at 1D_445256389, **(d)** shattering at 7B_703502819, **(e)** growth habit (TGH) at 1B_129176704, and **(f)** growth habit at 5B_586820792. Genotypes are grouped as homozygous reference, heterozygous and homozygous alternate classes, where present. The nucleotide substitution and Kruskal–Wallis *P* value are shown above each panel. Significant genotype-class differences were observed for TM at 5D_205488820 (*P* = 0.022), FLL at 6A_92352012 (*P* = 0.0452), SH at 1D_445256389 (*P* = 0.00361), SH at 7B_703502819 (*P* = 0.00561), TGH at 1B_129176704 (*P* = 0.00878), and TGH at 5B_586820792 (*P* = 0.00376), supporting the phenotypic effects of these GWAS-identified loci.

Shattering (SH) showed a highly significant major association on chromosome 1D and a secondary significant locus on chromosome 7B (Table 1). The chromosome 1D peak exceeded both BH-FDR and Bonferroni thresholds, and the QQ plots showed that most markers closely followed the null expectation except for the strongest associations. The most significant SNP was 1D_445256324 (P = 5.207 × 10⁻¹¹; BH-FDR = 1.128 × 10⁻⁶; Bonferroni-adjusted P = 1.128 × 10⁻⁶). Several neighboring SNPs, including 1D_445256334, 1D_445256342, 1D_445256363 and 1D_445256389, also showed very small P values, suggesting that these markers represent a shared LD block or a major QTL. A second significant association was identified at 7B_703502819 (P = 1.414 × 10⁻⁷; BH-FDR = 5.106 × 10⁻⁴; Bonferroni-adjusted P = 3.064 × 10⁻³). Genotype-class comparisons were consistent with the GWAS signals. At the representative 1D marker 1D_445256389, the Kruskal-Wallis test was significant (P = 0.00361). The C/C reference class showed shattering scores near 3.0, whereas the C/T class clustered near 1.0. A homozygous alternate class was not represented in the displayed data. At 7B_703502819, genotype classes also differed significantly (Kruskal-Wallis P = 0.00561); the C/C class showed the highest shattering scores, whereas the G/G and G/C classes remained lower. Together, the results indicate a major shattering-associated region on chromosome 1D with an additional contribution from chromosome 7B.

**Table 1.**
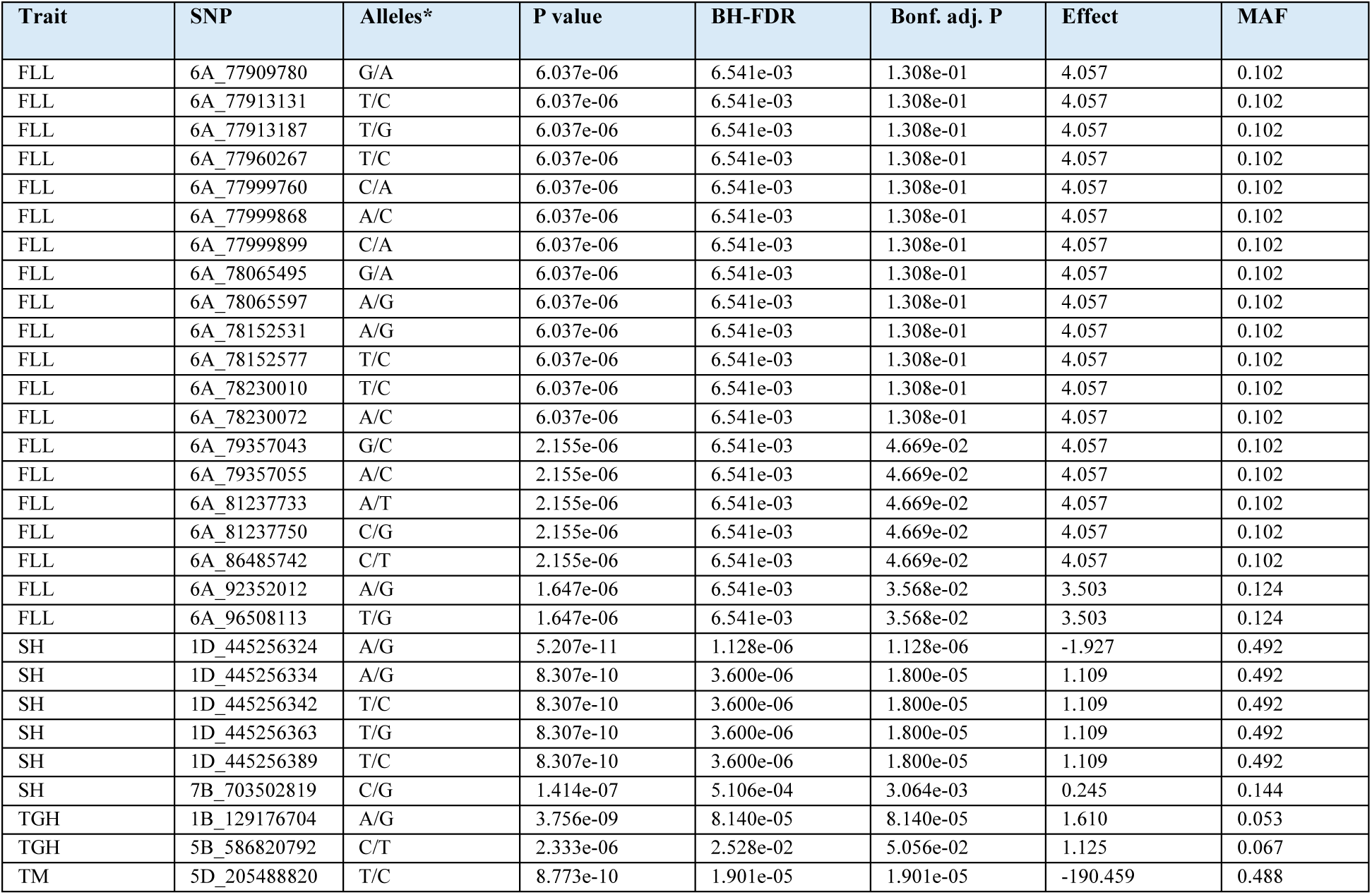
Significant marker-trait associations identified for flag leaf length, shattering, growth habit and tiller density in Pakistani wheat varieties. Significant SNPs detected by genome-wide association analysis are presented with their chromosomal position-based marker identifier, observed alleles, raw *P* value, Benjamini– Hochberg false discovery rate (BH-FDR), Bonferroni-adjusted *P* value, estimated allelic effect and minor allele frequency (MAF). Multiple closely linked SNPs on chromosome 6A were associated with flag leaf length (FLL), while shattering (SH) showed a major association block on chromosome 1D together with a secondary locus on chromosome 7B. Growth habit (TGH) was associated with loci on chromosomes 1B and 5B, whereas tiller density (TM) showed a strong association with 5D_205488820. Positive and negative effect values indicate the direction of the estimated allelic effect according to the allele coding used in the GWAS analysis. \***Alleles** are reported as the two nucleotide states observed at each SNP.

Flag leaf length (FLL) was associated primarily with a cluster of SNPs on chromosome 6A. Both BLINK and FarmCPU detected a pronounced 6A signal, and the QQ plots showed acceptable calibration, with departure from the expected distribution concentrated among the most significant markers. Several SNPs between approximately 77.9 and 78.2 Mb, including 6A_77909780, 6A_77913131, 6A_77913187, 6A_77960267, 6A_77999760, 6A_77999868, 6A_77999899, 6A_78065495, 6A_78065597, 6A_78152531, 6A_78152577, 6A_78230010 and 6A_78230072, shared essentially identical association statistics (P = 6.037 × 10⁻⁶; BH-FDR = 6.541 × 10⁻³) and a positive allelic effect of 4.057. The similarity among these statistics is consistent with strong LD among markers tagging the same underlying region. Additional SNPs at approximately 79.36, 81.24, 86.49, 92.35 and 96.51 Mb showed stronger evidence of association, and some passed Bonferroni correction. For the representative marker 6A_77909780, the Kruskal-Wallis test was significant (P = 0.0452). The G/G class showed a higher median FLL than the A/A class and the A/G class was intermediate. The direction of this descriptive trend was consistent with the positive allelic effect estimated by GWAS. Overall, the results support a chromosome 6A region associated with flag leaf length, but validation in larger populations and haplotype-based analyses will be important for refining the causal interval.

GWAS for tiller density (TM; tillers m⁻²) identified a major locus on chromosome 5D. Both BLINK and FarmCPU showed a distinct association peak in this region, and the QQ plots displayed upper-tail deviation consistent with a true association signal. The lead SNP, 5D_205488820, was highly significant (P = 8.773 × 10⁻¹⁰; BH-FDR = 1.901 × 10⁻⁵; Bonferroni-adjusted P = 1.901 × 10⁻⁵). The marker had a large negative estimated effect (−190.459 tillers m⁻²) and a MAF of 0.488, indicating that the associated polymorphism was common in the study panel. The genotype-class comparison was also significant (Kruskal-Wallis P = 0.022). The C/C reference genotype showed the highest tiller densities, with a median of approximately 320 tillers m⁻², whereas the C/T class showed substantially lower values, centred around 120-130 tillers m⁻². The T/T homozygous class was not represented in the displayed genotype-class plot. The direction of the phenotypic difference was consistent with the negative allelic effect estimated by GWAS. Taken together, these results identify 5D_205488820 as a strong candidate marker for tiller density that warrants independent validation.

## DISCUSSION

This study demonstrates that several morphologically defined DUS characteristics in Pakistani wheat germplasm have a detectable genetic basis and can be associated with discrete genomic regions. Significant marker-trait associations were identified for growth habit, shattering, flag leaf length and tiller density. These findings are relevant to DUS testing because conventional characterization remains predominantly phenotype-based, while the expression of some characteristics can be influenced by environment, developmental stage and observer-related variation. Recent studies have shown that historical DUS records can be integrated with genome-wide marker data to investigate the genetic architecture of DUS characteristics in wheat (Zanella *et al*., 2026), while genomic prediction has been proposed as a complementary tool for reference-collection management and trait-specific distinctness decisions (Roberts *et al*., 2026). The present study extends these approaches to Pakistani wheat germplasm and provides candidate genomic regions for future character-specific marker development.

Substantial phenotypic variation was evident across the evaluated DUS characteristics, as shown by the contrasting trait distributions and the separation of varieties into five broad phenotypic clusters. Most pairwise trait correlations were relatively weak, indicating that many descriptors captured distinct dimensions of morphology rather than redundant information. This is desirable for varietal characterization because highly correlated descriptors provide less independent discriminatory information. Similar analyses of wheat DUS datasets have also reported substantial variation in the diversity and genetic architecture of individual DUS characteristics (Zanella *et al*., 2026). The use of two complementary multi-locus GWAS methods further strengthened the analysis. FarmCPU iteratively combines fixed- and random-effect models to improve power while reducing confounding between tested markers and kinship (Liu *et al*., 2016), whereas BLINK improves computational efficiency and models associated loci without relying on arbitrary fixed LD bins (Huang *et al*., 2019).

Growth habit was associated most strongly with 1B_129176704, while a secondary signal was detected at 5B_586820792. The chromosome 5B association is biologically plausible because it lies in the broader genomic context of vernalization-related genes. Growth habit and seasonal adaptation are closely related, and winter wheat generally exhibits a more prostrate vegetative growth habit than spring wheat (Hyles *et al*., 2020). *VRN-B1* is a major MADS-box regulator of vernalization response and developmental transition in wheat (Milec *et al*., 2013). The chromosome 5B signal was reported to lie approximately 10 Mb from *VRN-B1* (TraesCS5B02G396600), placing it within the broad LD scale observed in the present germplasm. The homoeologous *VRN1* loci on chromosomes 5A, 5B and 5D are major determinants of seasonal growth habit and the transition from vegetative to reproductive development. Dominant *VRN1* alleles reduce or eliminate the vernalization requirement and promote spring growth habit, whereas recessive alleles at major *VRN1* loci are characteristic of winter growth habit (Santra *et al*., 2009). (Zhang *et al*., 2008) also demonstrated a strong relationship between allelic variation at *Vrn-A1*, *Vrn-B1*, *Vrn-D1* and *Vrn-B3* and variation in wheat growth habit. (Guedira *et al*., 2014) further showed that variation in the *VRN-B1* region affected vernalization requirement, winter dormancy release and heading date. Thus, the 5B association has both positional and functional plausibility. In contrast, the major 1B_129176704 signal could not be confidently assigned to a functionally validated wheat gene controlling vegetative growth habit based on the literature considered here. This locus may therefore represent a population-specific or previously unresolved region and should be prioritized for interval-level annotation and independent validation.

Flag leaf morphology is physiologically important because the flag leaf contributes substantially to assimilate production during grain filling, while its dimensions are quantitatively inherited and environmentally responsive (Wang *et al*., 2022). In the present study, FLL was associated with an extended set of markers on chromosome 6A. SNPs between approximately 77.9 and 78.2 Mb had nearly identical association statistics, indicating strong LD, while additional signals occurred at approximately 79.36, 81.24, 86.49, 92.35 and 96.51 Mb. This pattern could reflect one extended LD region or multiple linked loci affecting flag leaf development. Independent support is provided by (Zhang *et al*., 2026), who identified a leaf-length locus, QLfL_C&H-6A, spanning approximately 93.0-99.0 Mb and proposed TraesCS6A03G0284400 as a principal candidate gene. The present 96.51-Mb marker falls within that reported interval, and the 92.35-Mb marker lies immediately proximal to it. This physical correspondence strengthens the biological credibility of the distal 6A association. Haplotype analysis across the broader 6A region will be important for determining whether the 77-81-Mb and 92-97-Mb signals belong to a single extended haplotype or represent distinct loci. The lack of a significant Kruskal-Wallis test for 6A_77909780 also emphasizes that the broader locus or haplotype may be more informative than that individual SNP.

Shattering produced the strongest statistical association in the study, centred on 1D_445256324. The presence of several highly significant neighbouring SNPs around 445.26 Mb strongly suggests that these markers tag a common LD or haplotype block rather than multiple independent causal variants. A secondary association was detected at 7B_703502819, and genotype-class analyses supported phenotypic differentiation at both chromosome 1D and 7B loci. Shattering in wheat is closely related to rachis brittleness and spike disarticulation (Pourkheirandish and Komatsuda, 2022). Importantly, the present associations do not correspond to the canonical *BTR* loci responsible for the classical brittle-rachis domestication phenotype. The principal brittle-rachis loci are located on the short arms of homoeologous group-3 chromosomes; Br1/BTR-D occurs on 3DS, with related loci on 3AS and 3BS (Liu *et al*., 2022). Wheat *BTR1/BTR2* homologues have likewise been placed on chromosomes 3A and 3B (Akpinar *et al*., 2018). Consequently, the chromosome 1D association should not be annotated as *BTR1* or *BTR2*.

Rachis brittleness is, however, genetically more complex than the canonical *BTR* pathway alone. (Jiang *et al*., 2014) identified major brittle-rachis QTLs on chromosomes 2DS, 2DL, 3DS and 5AL in Tibetan semi-wild hexaploid wheat, demonstrating that loci outside chromosome group 3 can modify rachis disarticulation. The domestication gene *Q* on chromosome 5A provides another example: (Jiang *et al*., 2019) showed that a novel *Q* allele containing a 161-bp transposon insertion could generate brittle rachis through altered rachis cell-wall properties. (Peleg *et al*., 2011) also reported QTL affecting domestication-related spike characteristics, including a locus on chromosome 7B. Although older genetic maps do not allow confident physical colocalization with 7B_703502819, these studies support the existence of additional genetic modifiers of spike strength and disarticulation. The present 1D and 7B associations may therefore represent modifier loci affecting rachis strength, cell-wall composition, abscission-zone development or mechanical resistance. Because the 1D signal combines very strong statistical support with multiple neighbouring SNPs and clear genotype-phenotype differentiation, it is a high-priority target for conversion into a breeder-friendly assay and validation across independent DUS seasons.

Tiller density showed a strong association with 5D_205488820 that remained significant after both FDR and Bonferroni correction. The marker had a MAF of 0.488 and a large negative allelic effect, and the genotype-class comparison provided complementary phenotypic support. The involvement of chromosome 5D in wheat tillering has been reported previously. (Huang *et al*., 2003) identified tiller-number QTL on several chromosomes, including 5D, using advanced-backcross material. (Li *et al*., 2010) subsequently identified a major chromosome 5D QTL affecting tiller number at different developmental stages, and (Ren *et al*., 2018) identified the stable locus cqTN-5D.1 across multiple tillering stages. Collectively, these studies establish chromosome 5D as a recurrent genomic contributor to wheat tillering.

Nevertheless, previously reported 5D QTL cannot be assumed to be identical to 5D_205488820. (Ren *et al*., 2018) showed that 5D tillering QTL detected in different mapping populations were not necessarily the same locus, and older SSR-based studies lack the physical resolution required for direct correspondence with a SNP at 205.49 Mb. The present association therefore represents strong trait-and chromosome-level support for the importance of 5D, while the exact causal gene remains unresolved. Fine mapping of the LD-defined interval around 5D_205488820, followed by annotation of high-confidence genes and evaluation of their expression and functional relevance to branching pathways, would help identify plausible causal candidates. The genotype-count data underlying this locus should also be checked before publication because the reported MAF is high, whereas the displayed genotype-class plot does not contain a T/T homozygous class.

Taken together, the detected loci differ in the degree of external biological support. The distal chromosome 6A FLL region has particularly strong positional support because it overlaps an independently reported leaf-length interval. The chromosome 5D tiller-density signal is supported at the chromosome and trait level by several previous mapping studies. The 1D and 7B shattering loci are statistically strong but do not correspond to the classical group-3 *BTR* genes, while the major chromosome 1B growth-habit locus remains an unresolved candidate region. These differences emphasize the need to distinguish statistical association from causal-gene assignment and to validate each locus independently before regulatory or breeding application.

## IMPLICATIONS FOR MOLECULAR-ASSISTED DUS TESTING AND PLANT BREEDERS’ RIGHTS

The present findings have direct relevance to the emerging use of molecular information in DUS systems. (Zanella *et al*., 2026) demonstrated that loci associated with wheat DUS characteristics can be identified and that trait-associated SNPs can be converted into breeder-friendly KASP assays. Their work also showed that the predictive value of molecular markers varies substantially among traits. This distinction is important because markers intended to support DUS testing should remain demonstrably linked to the expression of recognized DUS characteristics rather than serving only as measures of overall genetic distance. (Roberts *et al*., 2026) similarly emphasized the complementary role of molecular information in phenotype-based DUS decisions. Their genomic-prediction framework uses historical DUS phenotypes and marker data to support reference-collection management and predict differences in individual DUS characteristics. The present study complements this approach by identifying discrete loci that may be suitable for a character-specific marker strategy.

For practical implementation in Pakistan, the chromosome 1D shattering region, chromosome 1B growth-habit locus, chromosome 5D tiller-density locus and distal chromosome 6A flag-leaf-length region represent priority targets for validation. Before any regulatory use, lead SNPs should be converted into low-cost assays such as KASP, tested in a larger independent collection of Pakistani varieties, and evaluated across the same environments and seasons used for official DUS characterization. Marker alleles should show repeatable correspondence with DUS notes, and their sensitivity, specificity, reproducibility and predictive error should be quantified. Haplotype-based assays may be more informative where several significant SNPs tag the same LD block, particularly for the chromosome 1D shattering and chromosome 6A flag-leaf-length regions. Molecular markers should therefore be regarded as complementary evidence that strengthens, rather than substitutes for, conventional multi-season DUS phenotyping.

### Conclusion

The present study provides a genome-wide assessment of the genetic basis of diagnostically important DUS characteristics in Pakistani wheat germplasm. The panel displayed substantial phenotypic diversity, extensive genetic admixture and hierarchical population structure, providing a suitable framework for association analysis. Robust genomic signals were detected for growth habit on chromosomes 1B and 5B, shattering on chromosomes 1D and 7B, flag leaf length on chromosome 6A and tiller density on chromosome 5D. Among these, the chromosome 1D shattering locus and chromosome 5D tiller-density locus showed particularly strong statistical support, while the distal chromosome 6A flag-leaf-length region also received independent positional support from published wheat studies. The major chromosome 1B growth-habit signal may represent a previously unresolved region and merits detailed functional investigation. These loci should be considered candidate regions rather than immediately deployable diagnostic markers. Independent validation across larger germplasm panels, environments and official DUS seasons is required, together with haplotype analysis, candidate-gene annotation and conversion of informative variants into breeder-friendly assays. If validated, character-specific molecular markers could improve the reproducibility and efficiency of varietal identification and provide an additional evidence layer for DUS decisions. Their most appropriate role is as a complement to conventional phenotyping, thereby strengthening the scientific basis of wheat variety protection and Plant Breeders’ Rights implementation in Pakistan.

## Contribution Statements

MA, RAJ and FS collected DUS characters data; IUH and MAH did GWAS and post-GWAS analysis; IAQ and ZA wrote manuscript; IUH and RAJ proof-read manuscript.

## Acknowledgement

Authors acknowledge the Dr. Awais Rasheed, Quaid-e-Azam University, Islamabad, Pakistan for providing genotyping assistance under the Science and Technology Partnership Program from the Ministry of Science and Technology (MoST) of China (KY202201001) and Higher Education Commission (HEC) NRPU grant number 15269.

